# Automated image segmentation uncovers the role of CD74^high^ human microglia in cognitive decline

**DOI:** 10.1101/2025.08.25.672200

**Authors:** Mariko Taga, Masashi Fujita, Neelang Parghi, Verena Haage, Andrew F. Teich, Julie A. Schneider, David A. Bennett, Ya Zhang, Philip L. De Jager

## Abstract

The role of activated microglia in Alzheimer’s disease (AD) is well established; the proportion of stage III activated microglia has been associated with AD and cognitive decline, but this morphologically defined subtype is relatively uncommon (1-2% of microglia) and its cellular function is unknown. Single-cell RNA-sequencing revealed CD74 as a marker gene that is enriched in immunologically active microglial subtypes associated with AD. Here, we evaluated the relationship between CD74 expression, AD-related traits, and microglial morphology using dorsolateral prefrontal cortex samples from two brain collections (ROSMAP: n=63, NYBB: n=91). An image segmentation pipeline using CellProfiler was developed to extract features from entire tissue sections. The pipeline automatically delineated gray and white matter regions and segmented 1,120,780 gray matter microglia. In a meta-analysis of the two datasets, we find an increase in frequency of microglia with high CD74 expression (CD74^high^) in relation to AD dementia (p = 0.038), particularly in the phase of terminal, accelerated cognitive decline before death. These microglia have a more rounded, amoeboid shape (ROSMAP: p = 1.4×10^-6^; NYBB: p = 2×10^-13^) which is a characteristic morphology of activated stage III microglia. Results were consistent across both datasets, highlighting the robustness of our cellular segmentation approach. This study identifies a potential role for CD74^high^ microglia and the CD74 ligand MIF in cognitive decline, and it provides evidence for a partially overlapping but distinct role for CD74^high^ microglia and morphologically defined stage III microglia, whose functional properties have remained poorly understood. These CD74^high^ microglia appear to be enriched for genes involved in cytokine response for class I and II antigen presentation, as well as regulation of T cell proliferation. These findings begin to link microglial subtypes defined by single-cell transcriptomic data with those characterized by classical morphological criteria to resolve the roles of different microglial functions to distinct stages in the trajectory to AD.

## Introduction

The heterogeneity of human microglia has become increasingly appreciated from both studies of freshly isolated microglia (from surgical resections or autopsy) and studies of nuclei extracted from frozen brain tissue^1–7^. One interesting subtype that has been defined transcriptomically in these single-cell/nucleus studies is characterized by high expression of MHC class II genes and genes involved in antigen presentation, such as CD74^2,4^. This microglial subtype has been associated with Alzheimer’s disease (AD)^2,4^ and displays expression of certain genes that are included in the genes defining the disease-associated microglia (DAM) 1 signature, but other microglial subtypes have higher expression of the DAM 2 signature^2,4^. While some studies have proposed this subtype as an intermediate state along a DAM trajectory^4,6^, it may represent a discrete state^4^ that is specialized in antigen presentation, a key function of microglia and other antigen presenting cells that serves to expand immune responses, including T cell proliferation.

While the role of T cells in humans with amyloid and tau proteinopathy remains unclear^8–10^, they seem to influence these proteinopathies in mouse models^8,11^. Single-cell and single-nucleus studies of microglia^1–4,7^ have clearly implicated certain subtypes in Alzheimer’s disease, which is defined pathologically by the presence of its two component proteinopathies, amyloid and tau. These association studies are fundamental in prioritizing certain subtypes of microglia for further evaluation; however, they do not inform us about the function of these subtypes and do not provide protein-based markers to facilitate further evaluations, such as examinations of their distribution and morphological characteristics in brain tissue. Transcriptomic studies do provide candidate genes for putative markers, but RNA-based hybridization methods such as RNAscope^12^ are difficult to scale to high throughput because of both cost and practical limitations due to current protocols. Here, we have developed an automated pipeline with which to scan and analyze tissue sections profiled by immunofluorescence with a key pool of markers that include an anti-CD74 antibody to capture the CD74^high^ microglial subtype that may represent an antigen presentating cell (APC)-specialized subtype in tissue sections from human neocortex. The entire tissue section is scanned, and the component images are segmented to identify the target cell types using CellProfiler (version 4.2.5)^13^. We demonstrated the robustness of the pipeline by capturing data from two different brain banks – the RUSH University AD Center and the New York Brain Bank – and combining results across the two sets of data that provide consistent results regarding the morphology of CD74^high^ microglia. We also assess the role of this microglial subtype in AD-related traits. Thanks to the automated, large-scale data capture we have also been able to assess large numbers of microglia. We focus on the dorsolateral prefrontal cortex as it is (1) a region affected by AD pathology, (2) affected later in the disease, avoiding the sclerosis that makes hippocampus and other regions more difficult to interpret in older individuals, and (3) deeply profiled by molecular profiles^3,14^. Finally, we leverage a new dataset of living microglia profiled with a CITE-seq panel to directly assess the transcriptomic profile of microglia defined as being CD74^high^.

## Materials and Methods

### Brain cohort

#### ROSMAP

The Religious Orders Study (ROS) and Memory and Aging Project (MAP) are both cohort studies conducted by the Rush Alzheimer’s Disease Center in Chicago^15,16^. Older individuals were free of known dementia at enrollment and agreed to annual clinical evaluation and brain donation at the time of death. Both studies were approved by an Institututional Review Board of Rush University Medical Center. All participants signed informed and repository consents and an Anatomic Gift Act.

Both cohorts conduct the same annual assessments, which include 19 cognitive functional tests, using validated procedures to diagnose AD and other dementias. All participants undergo a standardized structured assessment for AD which includes CERAD, Braak Stage, NIA-Reagan, and a global measure of AD pathologic burden with modified Bielschowsky. Amyloid load was measured by staining with three monoclonal antibodies against Aβ, which are 4G8, 6F/3D and 10D5, while PHF Tau tangles were measured using the AT8 antibody across eight brain regions. Both frozen and fixed brain tissues are accessible upon request from these subjects. AD pathology was assessed according to the National Institute on Aging-Reagan criteria . The clinical and pathologic methods have been previously reported^16,17^.

#### New York Brain Bank (NYBB)

The New York Brain Bank (NYBB) at Columbia University^18,19^ banked brain tissues coming from the Columbia Alzheimer’s Disease Research Center (ADRC), the NIA Alzheimer’s Disease Family Based Study (NIA-AD FBS) and, the Washington Heights, Inwood Columbia Aging Project (WHICAP). The Columbia ADRC cohort consists of clinical participants in the Columbia ADRC who have consent for brain donation. For all ADRC brain donations, one hemisphere is fixed while the other one is banked fresh and stored at -80°C. The NIA-AD FBS has recruited and followed 1,756 families from various ethnic groups with suspected late-onset AD, which include 9,682 family members and 1,096 unrelated, non-demented elderly. This initiative provides valuable biological resources such as brain tissue, plasma, and peripheral blood mononuclear cells (PBMCs) along with genetic data. The Resource Related Cooperative Agreement extended its recruitment efforts to include familial early-onset AD. Since 1992, the WHICAP has enrolled more than 6,000 participants, with representative proportions of African Americans (28%), Caribbean Hispanics (48%), and non-Hispanic whites (24%). Finally, the Biggs Institute Brain Bank at the University of Texas Health Science Center collects postmortem brain, spinal cord, cerebrospinal fluid, and dermal tissue from the study participants. The consent for the brain tissues was obtained from the donor’s legal next-of-kin before the autopsy. The left half hemisphere is fixed in 10% neutral-buffered formalin and the right hemisphere is banked fresh at -80°C. After a minimum of 4-week fixation period, the fixed tissue is sectioned and sampled in accordance with the NIA-AA 2012 guidelines AD neuropathologic guidelines.

### Immunohistochemistry

Six μm sections of formalin-fixed paraffin-embedded (FFPE) tissue from the dorsolateral prefrontal cortex (Brodmann Area 9) were de-paraffinized with CitriSolv (Xylene substitute, VWR) for 20 min. Heat-induced epitope retrieval was performed with citrate (pH=6) using a microwave (800W, 30% power setting) for 25 min. The sections were blocked with a blocking medium (3% BSA) for 30 minutes at room temperature (RT). The primary antibody anti-IBA1 (Wako; 011-27991; dilution 1:100) was incubated for 2 hours in RT followed by the secondary antibody incubation (monovalent to donkey/647 dye, 1/5000 dilution) for 1 hour. After washing with PBS, the sections were blocked again and incubated overnight at 4°C with antibodies against phosphorylated tau (AT8, Thermoscientific, MN1020B, dilution 1:500) and CD74 (Millipore Sigma, HPA010592, dilution 1:100). Tissue Sections were washed three times with PBS and incubated with fluorochrome-conjugated secondary antibodies (Thermo Fisher, dilution 1:500) for one hour at RT. After washing with PBS, GFAP-conjugated cy3 (Millipore Sigma, MAB3402, dilution 1:100) was incubated for 2H at RT. After three times washing, the sections were exposed to True Black Lipofuscin Autofluorescence Quencher to minimize the autofluorescence. Anti-fading reagent with DAPI (P36931, Life technology) was used for coverslipping.

### Image acquisition

The entire brain section was scanned for each subject using the Nikon Eclipse Ni-E fluorescence microscope. The tile-scanning imaging was performed twice: first at the low magnification (4X), and second at the high magnification (20X). The 4X imaging acquired the DAPI channel (emission wavelength: 460 nm, exposure: 100 ms) and the bright field channel (8 ms). The 20X imaging recorded five channels: DAPI (365 nm, 40 ms), pTau (488 nm, 40 ms), GFAP (565 nm, 40 ms), IBA1 (647 nm, 40 ms), and CD74 (750 nm, 40 ms). The image tiles of 4X were corrected for illumination unevenness, stitched into a whole-slide image using the Nikon NIS-Elements software, and saved as an image file in the ND2 file format.

### Tissue identification

Gray matter regions were identified from the low magnification images **(Fig. 1A)**. Both brightfield and DAPI channels of low magnification images were binarized by applying Otsu thresholding to each channel separately. Tissue regions were identified by taking the intersection of these binary images. Another Otsu thresholding was applied to the brightfield channel of the identified tissue to segment the gray matter region in the tissue.

**Fig. 1:**
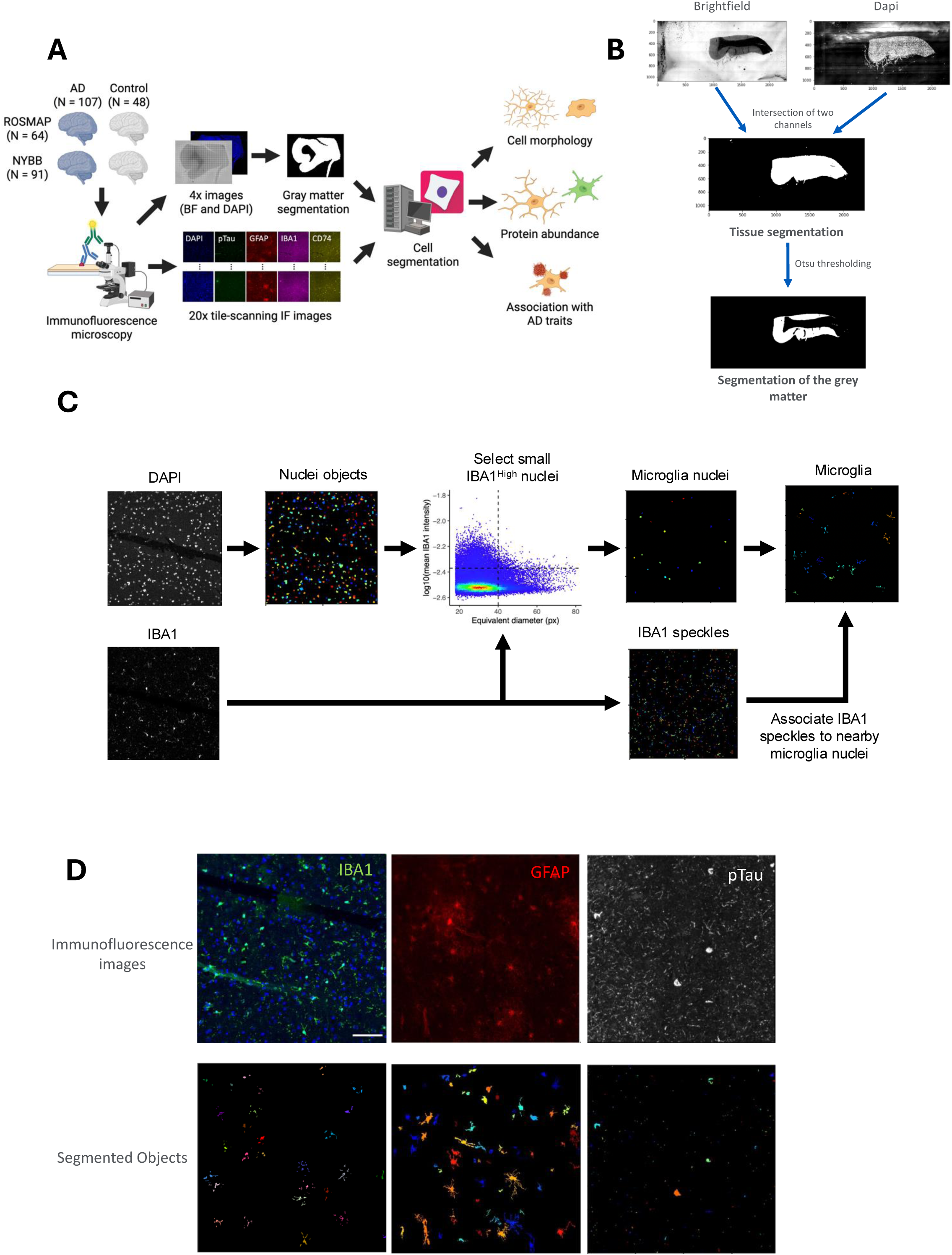
Study workflow and representative images. **(A)** The study workflow for image analysis, cell morphology measurement, and protein expression quantification using CellProfiler software. Paraffin-embedded brain tissues from ROSMAP (n=64) and NYBB (n=91) including AD (n=107) and control (n=48) brains were processed through tissue segmentation workflows to enable the measurement of cellular morphology and protein expression. Immunofluorescence staining was performed using antibodies against pTau, GFAP, IBA1, and CD74, with DAPI for the nuclear staining. Low-magnification (4X) brightfield and DAPI images were acquired for gray matter segmentation, followed by high-resolution (20X) tile-scanning IF imaging. Segmentation pipelines were applied to identify individual cells and quantify morphological features, protein abundance, and associations with AD-related traits. **(B)** Workflow for gray and white matter segmentation. The brightfield (4X) and DAPI (4X) images were combined, and the intersection of the two channels was used to define the tissue section. By applying Otsu thresholding, the gray matter was delineated from white matter using the CellProfiler software. **(C)** Illustration of microglial segmentation using CellProfiler software. Immunofluorescence images were acquired at 20X magnification, and IBA1 and DAPI channels were segmented. Nuclei from the DAPI staining that have smaller diameters and higher IBA1 intensities were selected as microglia nuclei. The threshold of IBA1 intensity was the mean + 2 standard deviation of IBA1 intensities. Cell bodies were defined by associating the speckles of IBA1 to microglia nuclei within a cutoff distance. Each color represents a distinct object. **(D)** Illustration of the final segmentation results for GFAP, IBA1, and pTau staining. (Left) IBA1 (green) and DAPI (blue). IBA1-positive objects associated with DAPI were segmented along with their cellular ramifications. (Middle) GFAP (red) -positive objects associated with DAPI were segmented including their extended processes. (Right) pTau (white) was segmented, including both neuropil threads and neurofibrillary tangles. Each color represents a distinct object. Scale bar, 100 μm.

### Quality check of image tiles

To perform a quality check of image tiles, the MeasureImageQuality module of CellProfiler was applied to all image tiles. The mean absolute deviation of IBA1 channels was visually inspected to exclude all empty and noisy image tiles. The blurry image tiles were detected through visual inspection of image tiles that were in the highest percentiles of the power log-log slope for IBA1, GFAP, and DAPI, based on the selected threshold. To detect image tiles with debris, the Otsu threshold of IBA1, GFAP, DAPI, pTau (AT8), and CD74 were applied, and the outliers were excluded.

### Image analysis and quantification

#### Nucleus

The image tiles that passed the quality check were processed through the automated image analysis pipeline of CellProfiler software 4.2.5 version. Nuclei were identified and segmented by applying the ’IdentifyPrimaryObjects’ module to the DAPI channel. The advanced settings were applied to detect objects with diameters ranging between 18 and 80 pixels. All objects with diameters outside this range were excluded, and the “Robust Background” method was used for thresholding. The “Shape” method was used to differentiate clumped objects and to delineate dividing lines between them (**Fig. 1B**).

#### Microglia

Microglia objects were segmented in a two-step approach, (i) selecting microglia nuclei from all nuclei objects, and (ii) attaching cytoplasmic areas to each microglia nuclei. Microglia nuclei were selected so that they have equivalent diameter < 40 pixels (13.0 μm) and their IBA1 fluorescent intensities are higher than the mean and 2 standard deviation in all nuclei. To segment microglia cytoplasm, image enhancement was applied to the IBA1 channel (feature type, ‘Neurites’; enhancement method, ‘Tubeness’; smoothing scale, 4). Fragments of microglia cytoplasm were identified by applying the ’IdentifyPrimaryObjects’ module to the enhanced IBA1 images (typical diameter, 10–40 pixels; thresholding, ‘Robust Background’; declumping, ‘Intensity’). Each fragment of microglia cytoplasm was assigned to the nearest microglia nuclei if they are within 60 pixels (19.5 μm).

#### Astrocytes

First, to maximize the detection of astrocyte ramifications, the “EnhanceOrSuppressFeatures” function was utilized. The “Enhance” option was selected with the enhancement method set to “Line structures” and the feature type specified as “Neurites.” The feature size, defined as the diameter of the largest speckle, was set to 50. Then, GFAP+ objects were identified and segmented using the ’IdentifyPrimaryObjects’ module. The advanced settings were adjusted to detect objects ranging from 10 to 300 pixels in diameter. The ’SplitOrMergeObjects’ module was utilized to merge small objects, such as ramified structures, and associate them with nearby larger objects. The object resulting from this merging process was named “GFAPmerged.” Once again, the “EnhanceOrSuppressFeatures” module was employed to enhance the small ramified objects of astrocytes; however, this time, the enhancement method “Tubeness” was applied, and the smoothing scale was set to 2.0. The image resulting from this image processing was named “GFAPTubeness.” Using the “IdentifiySecondaryObjects” module, “GFAPmerged” objects were associated with the “GFAPTubeness” image using ‘propagation” as the method of identifying the secondary objects. The adaptative threshold strategy was used with the Robust Background thresholding method. The averaging method applied was “Mean,” while the variance was calculated using the standard deviation method. The “SplitOrMergeObjects” module was used to merge all ramifications and assign them to the nearest largest object, resulting in the creation of a new object named “GFAP_final.” Finally, using the “RelateObjects” module, with nuclei as the parent objects and “GFAP _final” as the child objects, we filtered IBA1+DAPI+ objects. The intensity, as well as size and shape, were measured using the “MeasureObjectSizeShape” and “MeasureObjectIntensity” modules.

<u>pTau (AT8):</u> To segment pTau objects, “IdentifyPrimaryObjects” module was applied to the image tiles of pTau channel with typical diameter 1–80, robust background thresholding, and 20 standard deviation as a threshold. Segmented pTau objects were further classified using R: objects with areas less than 900 pixels were labeled as “pTau threads”, and objects with areas above 900 pixels and below 10,000 pixels were classified as” pTau tangles”.

### Human microglia isolation for CITE-seq

Autopsy samples of human brains were obtained from Rush University Medical Center/Rush Alzheimer’s Disease Center (RADC) in Chicago, IL (Dr. Bennett) and Columbia University Medical Center/New York Brain Bank in New York, NY (Drs. Vonsattel and Teich), as well as surgically resected brain specimens from Brigham and Women’s Hospital in Boston, MA (Drs. Sarkis, Cosgrove, Helgager, Golden, and Pennell), Rocky Mountain Multiple Sclerosis Center, Denver, CO (Dr. John Corboy) and Alzheimer’s Disease Research Center/Precision Neuropathology Core of University of Washington School of Medicine (Dr. C. Dirk Keene).

All brain specimens were collected through informed consent and/ or brain donation programs at the respective organizations. All procedures and research protocols were approved by the corresponding ethical committees of our collaborator’s institutions as well as the Institutional Review Board (IRB) of Columbia University Medical Center (protocol AAAR4962).

Once the tissue samples were received, they were weighed, and up to 2g of tissue was used for microglia isolation. The pH of all buffers was adjusted to 7.3-7.4 and stored at 4°C until use. A dounce homogenizer (Carl Roth, Cat#: CXE2.1) was placed on ice and filled with 12mL of ice-cold Hibernate-A (Thermo Fisher Scientific; Cat#: A1247501). The tissue was minced in a pre-chilled petri dish with 3-4 mL Hibernate-A (Thermo Fisher Scientific; Cat#: A1247501), then transferred to the dounce homogenizer placed on ice. The minced tissue was then dissociated using the loose pestle and subsequently, the suspension was passed through a prewet 100µM strainer (Corning, Cat#: CLS352360) into a 50mL falcon tube (fisher scientific, Cat#: 10203001). The single-cell suspension was centrifuged at 400g, for 10 min at 4°C. Meanwhile, isotonic percoll was prepared by adding 1mL 10x DPBS (pH = 7.2-7.4; Thermo Fisher Scientific; Cat#: 14200075) to 9mL pH-adjusted percoll solution (Fisher Scientific; Cat#: 10607095). Following centrifugation, the supernatant was aspirated and DPBS was added to reach a final volume of 12mL. Next, 4mL of isotonic percoll was added, and the solution was thoroughly mixed. Then, the solution was overlayed with 16mL of DPBS (Corning; Cat#:21-040-CV) by slightly tilting the falcon tube and keeping the pipetboy tip against the tube wall to avoid mixing the layers. The layered sample was centrifuged at 3000g for 10 min at 4°C and the supernatant, including the myelin disk in between the two phases was aspirated. 15mL of cold DBPS (Corning; Cat#:21-040-CV) was added for washing, and the falcon tube was tilted three times at 180°C and subsequently centrifuged at 400g for 10 min at 4°C. To isolate microglia from the myelin-depleted cell suspension, cells were first counted using a Nexcelom cell Counter, followed by Magnetic-Activated-Cell-Sorting (MACS) technology following the manufacturer’s instructions.

Briefly, the cell suspension was centrifuged at 300g, 4°C for 10 min, then the supernatant was completely removed. The cell pellet was resuspended in 80 µl of MACS buffer for every 10^7^ cells. For every 10^7^ total cells, 20µl of CD11b MicroBeads (Miltenyi; Cat#: 130- 093-636) were added, thoroughly mixed, and incubated for 15 min at 4-8°C. The cells were then washed by adding 1-2 mL of buffer per buffer per 10^7^ total cells and centrifuged at 300g for 10 min at 4°C. The supernatant was completely removed, and up to 10^8^ cells were resuspended in 500µl of MACS buffer (PBS (Corning; Cat#: 21-040-CV), 0.5% BSA (pluriSelect; Cat#: 60-00020-10BSA), 2mM EDTA (Thermo Fisher Scientific; Cat#: AM9260G)).

A magnetic column (Miltenyi, LS coumns, Cat#: 130-042-401) was placed into the magnetic field of a suitable MACS separator (Miltenyi, Cat#: 130-042-303) and prepared by rinsing with 3mL of MACS buffer. Subsequently, the cell suspension was applied to the column, followed by three washing steps by adding 3mL of MACS buffer. The column was then removed from the separator, and placed on a 15mL falcon tube (Corning, Cat#: CLS430790) for collection. Then, 5mL of MACS buffer was added, and magnetically labeled cells were flushed. Isolated CD11b positive cells were centrifuged at 300g for 10min at 4°C and resuspended in 1-2 mL of DPBS (Corning; Cat#:21-040-CV) for cell counting.

### CITE-Seq staining

After cell counting, cell viability was assessed, and 0.7×10^6^ cells were transferred to a 5mL low-protein binding tube (Fisher Scientific; Cat#: 13-864-407). 1mL of cold PBS (Corning; Cat#:21-040-CV) was added, and the cells were filtered through a blue lid filter (FACS tube) back into the 5ml low-protein binding tube (fisher scientific; Cat#: 13-864- 407). The sample was then centrifuged at 300g for 10 min at 4°C, with the washing step repeated. The supernatant was removed, and the cell pellet was resuspended in 22.5µL of Cell Staining Buffer (BioLegend, Cat#: 420201) and 2.5µL of Human FcBlock (BioLegend, Cat#: 422301) was added for 10 min incubation on ice.

The hashing master mix was prepared as follows: 2µL of TS-A hashtag antibody (0.5 mg/mL, BioLegend, TotalSeq- A0251 anti-human Hashtag1, Cat#: B364030; TotalSeq- A0252 anti-human Hashtag2, Cat#: B369100; TotalSeq- A0253 anti human Hashtag3, Cat#: B378300; TotalSeq- A0254 anti human Hashtag4, Cat#: B381073) was added to 20µL of Cell Staining Buffer (BioLegend, Cat#: 420201) and passed through a 0.1µm filter at 12000g for 2 min at 4°C. Next, 6.25µl of reconstituted combined CITE-Seq cocktail was added to 18.75µl of hashtag master mix. The mixed solution was added to the cells incubating in the FcBlock and Cell Staining Buffer.

If samples were not hashed, 6.25µl of reconstituted combined CITE-Seq cocktail was added to 18.75µl of Cell Staining Buffer. The mixture was then added to the cells and incubated in FcBlock and Cell Staining Buffer. After 30 min of incubation, four washing steps were performed, each consisting of adding 1mL of cells of Cell Staining Buffer followed by a centrifugation at 300g for 10min at 4°C. Before the final wash, the cells were filtered through a blue lid filter cap. Cells were then counted and processed further for single-cell sequencing.

### Single-cell RNA sequencing with CITE-seq

Detailed single-cell RNA sequencing procedures were performed following the Biolegend TotalSeq™-A Antibodies and Cell Hashing with 10x Single Cell 3’ Reagent Kit v3.1 (Dual Index) protocol (https://www.biolegend.com/fr-ch/protocols/totalseq-a-dual-index-protocol) with minor modifications. Briefly, TotalSeq™-A antibody-labeled and hashed microglia cells from different brain regions of the same donor were pooled and then counted using a Nexcelom Cellometer Vision with a 10x objective and AO/PI stain. Single-cell library generation was constructed using 10x Chromium Next GEM Single Cell 3’ Reagent Kits v3.1 (Dual Index) with Feature Barcode technology for Cell Surface Protein (10x Genomics, Pleasanton, CA) according to the manufacturer’s protocol with modification. Briefly, a total of 10,000 – 20,000 cells were loaded into one well of the Chip G kit for GEM generation using a 10x Genomics chromium controller single-cell instrument.

Incubation of the GEM generated barcoded cDNA from polyadenylated mRNA and simultaneously generated DNA from the cell surface protein Feature Barcode within the same single cell in the GEM. ADT additive primer (0.2µM Stock) and HTO additive primer v2 (0.2µM Stock) were used to generate feature barcoded-DNA during complementary DNA (cDNA) amplification to enrich TotalSeq-A cell surface oligonucleotides. Cell barcoded cDNA molecules were amplified by PCR and used to construct libraries. The amplified cDNA was then separated by SPRI size selection into cDNA fractions containing mRNA derived cDNA (>400bp) and ADT/HTO-derived cDNAs (<180bp) and further purified by additional rounds of SPRI selection. During sample index amplification, independent sequencing libraries were generated from the mRNA, ADT and HTO cDNA fractions using Dual-Index Kit TT, Set A (PN-1000215), Dual Index ADT (DI_ADTx) i5/i7 Primer Pairs (10µM Stock) and Dual Index HTO (DI_HTOx) i5/i7 Primer Pairs (10µM Stock). These libraries were analyzed and quantified using TapeStation D5000 screening tapes (Agilent, Santa Clara, CA) and Qubit HS DNA quantification kit (Thermo Fisher Scientific). The libraries were pooled and sequenced on a NovaSeq 6000 with S4 flow cell (Illumina, San Diego, CA) at a ratio of GEX:ADT:HTO = 8:4:1. Paired-end, dual-index sequencing was used to perform 28 cycles for read 1, 10 cycles for i7 index, 10 cycles for i5 index, and 90 cycles for read 2.

### CITE-Seq data analysis

The raw sequencing data for each sample were processed using the Cell Ranger software (v.7.2.0) from 10x Genomics. This was conducted on an HPC cluster configured with 24 GB of memory and 4 local cores. The GRCh38 human reference transcriptome was used (also provided by 10x Genomics). A custom feature reference was utilized for the detection of unique molecular identifiers (UMIs) and enabling precise feature barcoding. Further analysis was carried out using Seurat v5.0.3 in Rstudio v4.3.1. Separate matrices for gene expression and antibody capture (ADT) were integrated into a Seurat object for comprehensive analysis. Sum and median UMI counts across cells were calculated for each antibody. These counts were used to create visualizations of the UMI distribution, using log-scale transformations. Quality control measures included ADT normalization using the CLR transformation method.

Among the nine libraries of human microglia, we selected the PV053 library because it had the best quality. We defined microglia as CD74^high^ if its CD74 protein abundance was higher than mean + 2 standard deviations of CD74 protein abundance in the library. The differential expression analysis of proteins and RNA was performed using FindMarkers() function of the R package Seurat.

### Terminology

*Tangles_mf:* Tangles identified by immunohistochemistry using the anti-AT8 antibody in mid frontal cortex.

*Tangles:* Tangle score in 8 brain regions (Hippocampus, entorhinal cortex, mid frontal cortex, inferior temporal, angular gyrus, calcarine cortex, anterior cingulate cortex, and superior frontal cortex).

*NFT_mf*: Neurofibrillary tangles burden measured by sliver staining in mid frontal cortex. *NFT:* Neurofibrillary tangles burden measured by sliver staining in 5 brain regions (mid frontal cortex, mid temporal cortex, inferior parietal cortex, entorhinal cortex, and hippocampus).

*Braaksc:* A semiquantitative measure of NFT pathology severity. Bielschowsky silver stain was used to visualize NFTs in the frontal, temporal, parietal, entorhinal cortex, and hippocampus.

## Results

### Developing an automated pipeline capable of segmenting microglia and astrocytes in a human brain tissue section

To investigate the distribution of CD74^high^ microglia across a large number of donors, we co-stained fixed tissue samples from the dorsolateral prefrontal cortex (DLPFC) with antibodies targeting CD74, IBA1 (microglial marker), GFAP (astrocytic marker), and phosphorylated Tau (pTau, AT8 antibody) (see Methods). This specific brain region was selected as it is affected by AD pathology and has been the substrate for many rounds of “omic” data generation of samples from the Religious Order Study and Memory & Aging Project (ROSMAP)^3,4,14,20,21^. To capture many of microglia and astrocytes from each individual, the entire tissue section was scanned using a Nikon ECLIPSE Ni immunofluorescence microscope with an automated stage at 4X for tissue identification purposes and at 20X resolution for image capture. The 20X resolution was selected to balance the ability to run samples in a moderate-throughput manner with the acquisition of images of sufficient resolution to capture the proximal ramification of microglia. Each sample took approximately 2-3 hours to scan, depending on the tissue section size. A median of 130 tiles was captured per tissue section in the ROSMAP samples. We also deployed the same experimental pipeline to profile fixed tissue samples from the same region collected from donors undergoing brain banking at the New York Brain Bank (NYBB)^19^; these tissue sections were larger, so we focused image capture and segmentation in the gray matter for this collection of samples.

The image segmentation pipeline first involves using both images of the tissue stained with 4′,6-diamidino-2-phenylindole (DAPI) (which identiies nuclei) and brightfield 4X images to separate the gray from the white matter. Individual image tiles were used in subsequent analysis if more than 95% of its area belonged to gray matter (see Methods)(**Fig. 1A**). These gray matter images were then processed using CellProfiler software version 4.2.5. Our pipeline for microglial segmentation begins with the identification of nuclei in each image tile using the DAPI stain. Next, nuclei were filtered by their sizes and IBA1 intensities to identify microglial nuclei (see Methods for details). To detect the ramifications of microglia, small IBA1 blobs were linked to nearby microglial nuclei and assembled into microglia objects using the SplitOrMergeObjects and RelateObjects features of CellProfiler (**Fig. 1B**). Segmentation of astrocyte objects was performed by identifying GFAP+ objects that overlap with DAPI+ objects (nuclei) and filtering them by size (**Fig. 1C**). Threads and tangles of phosphorylated tau (pTau) were segmented as primary AT8+ objects regardless of the presence of DAPI (**Fig 1C**). Following object segmentation, CD74 and IBA1 mean fluorescence intensities were measured within the microglia objects, and GFAP intensity was measured within the astrocyte objects. CD74^high^ microglia were defined among microglia objects as those microglia which had an expression of CD74 greater than the mean of CD74 expression plus two standard deviations. The morphology of each segmented object is measured, providing more than 30 different metrics, including the volume, area, perimeter, and compactness. Individual objects are aggregated per tile and then per section to produce a count of each type of object for each participant. The X/Y coordinates of each object are also recorded, allowing us to detect their distribution in the tissue sample. We applied this staining protocol and automated segmentation pipeline to 64 participants from the ROSMAP studies and 91 donors from the NYBB, totaling 155 individual brain samples.

Detailed demographic, neuropathologic, and clinical characteristics for these two sets of samples are provided in **Table. 1**. Due to variability in tissue quality, we observed higher background staining levels in the NYBB samples (see **Supplementary Fig. 1**). As a result, image analysis was conducted separately for the two brain banks. However, the automated image segmentation pipeline performed well in both sets of samples. A total of 157,120 microglia were segmented in ROSMAP and 326,613 microglia in NYBB, which provided larger tissue samples. This resulted in the same median density of 48 microglia per mm² of gray matter for both sample sets. To validate the reliability of our automated pipeline for quantifying microglia, we selected 40 ROSMAP samples that have both immunofluorescence (IF)-derived data from this study and single-nucleus RNA-seq data generated in an earlier study^3^ from the DLPFC of the opposite hemisphere. We observed a positive Pearson’s correlation coefficient in the microglial proportions derived from IF images (denominator being the total number of DAPI+ objects) and single-nucleus RNA-seq data (denominator being all sequenced nuclei) (r=0.49, p=0.0013) (**Fig. 2A**). A positive correlation was also observed between the proportion of GFAP+ astrocytes derived from IF images and the expression level of GFAP in bulk RNA-seq data (r=0.35, p=0.0053) in the 63 ROSMAP participants and in bulk proteomics (r=0.48, p=0.0057) in the 32 ROSMAP participants (**Fig. 2B**), further supporting the robustness of the segmentation pipeline.

**Table. 1:** Demographic table for the ROSMAP and NYBB cohorts.

|  | ROSMAP | NYBB |
| --- | --- | --- |
| Number of participants | 64 | 91 |
| Male (%) | 32 (50%) | 33 (32%) |
| Age at death, median (Q1 – Q3) | 88 (83 – 92) | 88 (79 – 90) |
| Pathological AD (%) | 32 (50%) | 75 (83%) |
| Clinical diagnosis (%) | AD, 18 (28%) | Not Available |
|  | MCI, 24 (38%) |  |
|  | NCI, 22 (34%) |  |

**Fig. 2:**
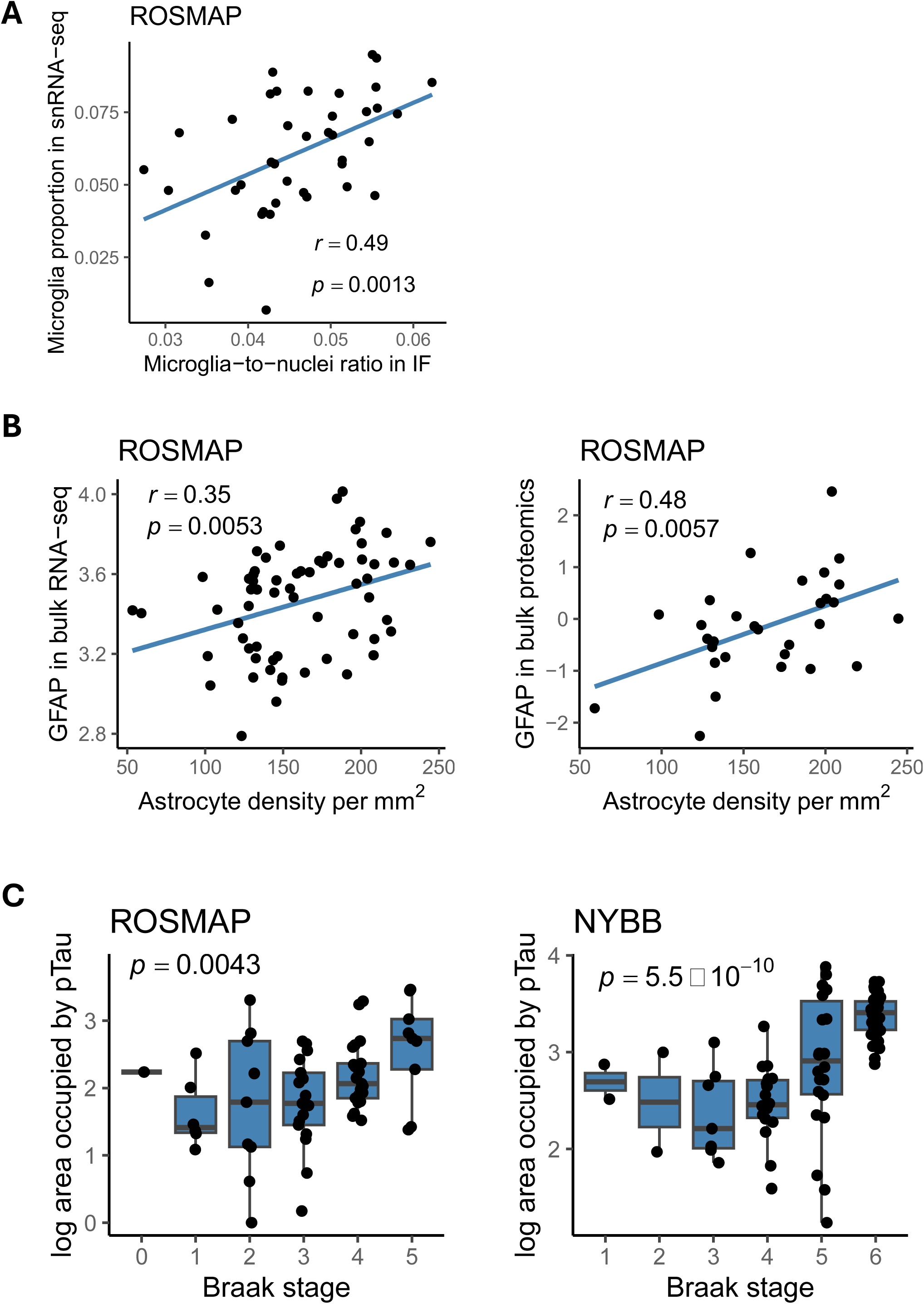
Evaluation of automated segmentation performance. **(A)** Scatter plot showing the correlation between microglia proportions from the snRNA-seq data (y-axis) and the microglia-to-nuclei ratio from the immunofluorescence staining (x-axis) in the same individuals from the ROSMAP cohort (n=40). A positive correlation was observed (Pearson’s correlation coefficient, r = 0.49; p = 0.0013). The linear regression line is shown, with the shaded area representing the 95% confidence interval. Each dot represents an individual sample. **(B)** Scatter plots showing positive correlations between GFAP+ cell density per mm² from immunofluorescence staining (x-axis) and GFAP levels measured by bulk RNA-seq (y-axis; Pearson’s r = 0.35, p = 0.0053) and proteomics (y- axis; Pearson’s r = 0.48, p = 0.0057) in the same individuals from the ROSMAP cohort. Each dot represents an individual sample. **(C)** Boxplots showing the accumulation of pTau increasing with Braak stage in both ROSMAP and NYBB cohorts. The area occupied by ptau (AT8) obtained from IF data increases across Braak stages in the ROSMAP (left, *p*=0.0043) and NYBB (right, *p*=5.5×10-^10^). Center line: median; box limits: upper and lower quartiles; whiskers: the most extreme data point which is not more than 1.5 times the inerquartile range from the box. Each dot represents an individual sample.

For pTau quantification, we used the “Area occupied” by pTau (AT8 antibody) as measured by CellProfiler. In both ROSMAP and NYBB cohorts, the area occupied by pTau were significantly and positively associated with Braak stage and a pathological diagnosis of AD (**Fig. 2C** and **Supplementary Fig. 2B**). In the ROSMAP samples, where quantitative pathological measurements were available, the area occupied by pTau also showed significant positive associations with several measures derived from the fixed hemisphere include a measure of AT8 immunohistochemistry staining in the midfrontal cortex, the average AT8 immunostaining across eight brain regions, a measure of neurofibrillary tangles derived from a silver stained images of the midfrontal cortex, and the average neurofibrillary tangle burden from silver stained images of five brain regions (seeTerminology)(https://www.radc.rush.edu/docs/var/overview.htm?category=Pathology) (**Supplementary Fig. 2B**). Additionally, when differentiating between pTau threads and tangles in our images based on their sizes (see Methods), we found that both have a positive association with Braak stage (**Supplementary Fig. 2A**). However, the burden of pTau threads showed stronger associations with pathological traits than that of pTau tangles (**Supplementary Fig. 2B**).

### Microglial density is similar in AD and non-AD brains

We compared microglial density between participants with and without a pathologic diagnosis of AD in ROSMAP and NYBB (**Fig. 3A**). In ROSMAP, non-AD participants have a median microglia density of 48.3 (Q1 – Q3: 42.3 – 52.9) per mm², while those with AD have a density of 46.8 (43.1 – 54.0) per mm². In NYBB, non-AD participants have a median density of 41.7 (37.3 – 50.1) per mm², and participants with AD have a density of 49.1 (40.8 – 57.2) per mm². However, these differences are not significant: p = 0.66 for ROSMAP and p = 0.24 for NYBB. In ROSMAP, a participant’s ante-mortem status is also available. The density of microglia showed no significant differences among those with no cognitive impairment (NCI), mild cognitive impairment (MCI), and AD dementia (p = 0.13) (**Fig. 3B**).

**Fig. 3:**
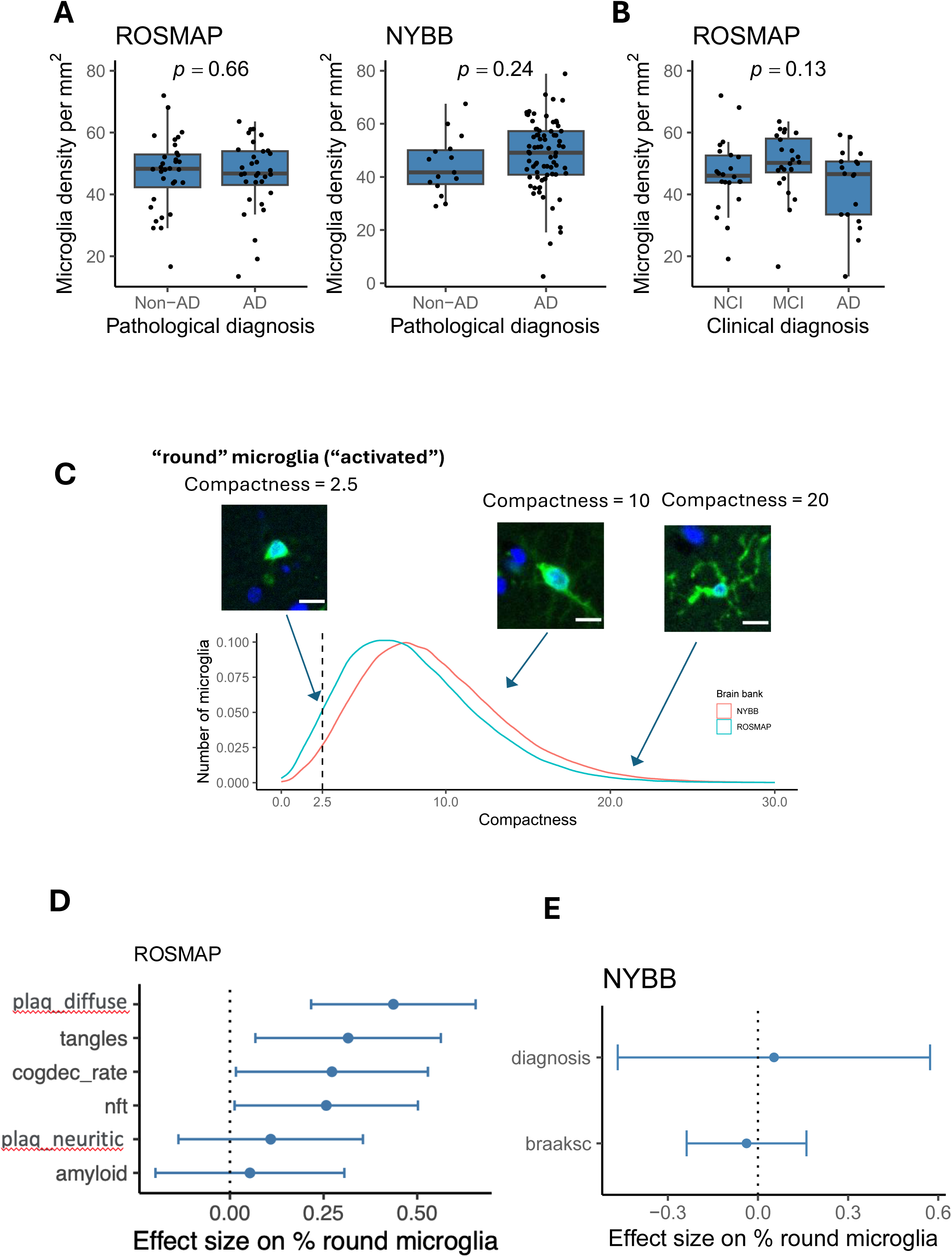
Morphological characterization of rounded microglia in AD pathology. **(A)** Boxplots showing IBA1⁺ cell density (cells/mm²) associated with pathological diagnosis in ROSMAP and NYBB cohorts. Microglia density was assessed based on IBA1 immunostaining. Each dot represents an individual sample. **(B)** Boxplots showing IBA1⁺ cell density (cells/mm²) associated clinical diagnoses in ROSMAP cohort. No significant association was observed (*p* = 0.13). Each dot represents an individual sample. Center line: median; box limits: upper and lower quartiles; whiskers: the most extreme data point which is not more than 1.5 times the interquartile range from the box. The *p*-values were computed using *t*-tests within linear models, adjusting for age, sex, dichotomized background staining level, and, for ROSMAP samples, post-mortem interval. **(C)** Distribution of microglial morphological compactness in ROSMAP and NYBB cohorts. The density plots shows the distribution of microglial compactness scores in the NYBB (red) and ROSMAP (blue) cohorts. The dotted line shows the threshold for dichotomizing microglia into “rounded” and “non-rounded”. Representative immunofluorescence images of IBA1 (green) and DAPI (blue) illustrate examples of microglia with compactness 2.5 (low), 10.0 (intermediate), and 20.0 (high). Scale bars= 10 μm. **(D, E)** Association between the proportion of rounded microglia and AD pathological and clinical traits in ROSMAP and NYBB cohorts. The dots show the effect size of each pathological trait on the proportion of rounded microglia, estimated using a linear model adjusted for age, sex, and dichotomized background staining level; post-mortem interval was additionally adjusted for in the ROSMAP samples. Error bars indicate 95% confidence intervals.

### Morphologically rounder microglia are associated with AD pathology

We assessed the morphological characteristics of microglia that we segmented in the ROSMAP and NYBB samples focusing on their roundedness, as activated stage III microglia are traditionally characterized as rounded, dense cells with retracted processes^20,22^. Leveraging the morphological measures of IBA1+ cells obtained by CellProfiler, we prioritized the evaluation of the “compactness” measure, defined as the mean squared distance of an object’s pixels from its centroid, divided by its area. A filled circle has a compactness of 1, while irregular objects or branched shapes have a higher compactness (greater than 1). Compactness was selected over eccentricity measurement to assess microglial morphology, as eccentricity is more suitable for distinguishing round versus elongated shapes and is less informative for evaluating cellular complexity or branching structures.

We first created an operational definition to identify the extreme end of the microglial morphological spectrum: we defined “rounded” IBA1+ microglia as cells with a compactness value of 2.5 or below (**Fig. 3C**). As shown in **Fig. 3D**, in the ROSMAP sample, the percentage of round microglia showed a positive association with diffuse amyloid plaques (p=1.96×10^-4^), tangles (p=0.0134), neurofibrillary tangles (p=0.0398), and rate of cognitive decline (p=0.0378), illustrating the increased proportion of activated microglia found with worsening AD-related traits, consistent with prior reports that used manual counts of stage III microglia^20,22^. The NYBB samples do not have quantitative measures of pathology, so we evaluated the percentage of round microglia in those samples in relation to Braak stage (p = 0.71) and a pathological diagnosis of AD (p = 0.84), measures that are not as sensitive as those used in ROSMAP (**Fig. 3E**).

### CD74^high^ microglia are more frequent among participants with cognitive decline

A small fraction of microglial cells showed high levels of CD74 protein expression (**Fig. 4A**). We defined microglia as CD74^high^ when their CD74 mean fluorescence intensity was higher than the mean plus 2 standard deviations of all microglia in the same sample (**Fig. 4B**). Next, we assessed the association of CD74^high^ microglia with AD (**Fig. 4C**). There was no significant difference in the proportion of CD74^high^ microglia between subjects with and without a pathologic diagnosis of AD in both ROSMAP (p=0.91) and NYBB (p=0.39). Additionally, no association was found between Braak stage and the proportion of CD74^high^ microglia in both sample collections (p=0.63 for ROSMAP and p=0.40 for NYBB), suggesting that the proportion of CD74^high^ microglia is not associated with the presence of AD pathology (**Supplementary Fig. 3**).

**Fig. 4:**
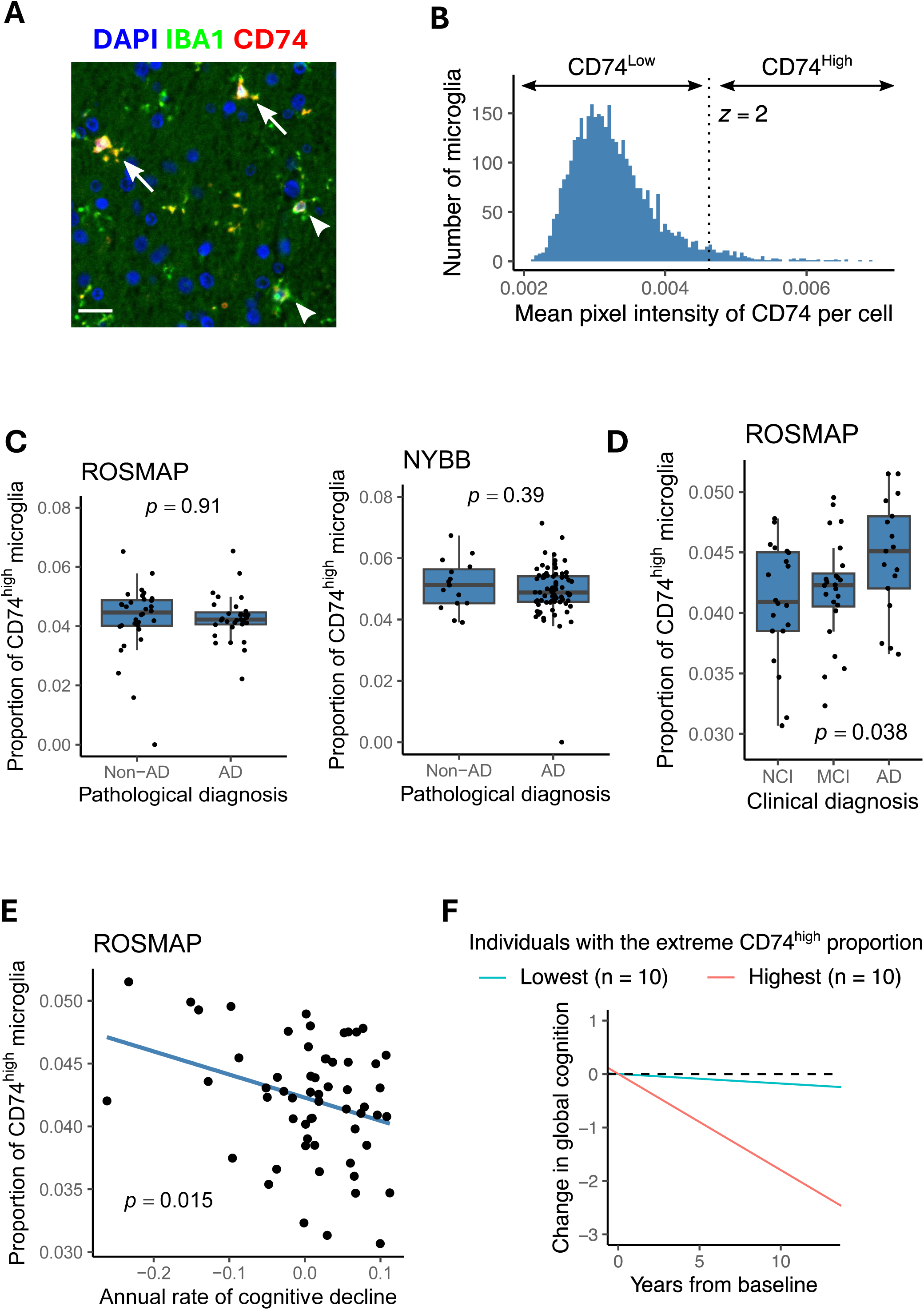
Characterization of CD74^high^ microglia and their association with AD traits. **(A)** Illustration of CD74^high^ microglia (arrows) and CD74^low^ (arrowheads) stained with IBA1 (green) CD74 (red), and DAPI (blue). Scale bar= 20μm. **(B)** The histogram shows the distribution of microglial CD74 intensity in the ROSMAP samples. The dotted line shows the mean + 2 standard deviations of CD74, used as the threshold for dichotomizing microglia into CD74^high^ and CD74^low^ subsets. **(C)** Box plots illustrating the association between the proportion of CD74^high^ and the pathological diagnosis of AD in the ROSMAP (left) and NYBB (right) cohorts. Each dot represents an individual sample. **(D)** Box plots illustrating the association between the proportion of CD74^high^ and the clinical diagnosis of AD in the ROSMAP samples. Each dot represents an individual sample. ***(C,D)*** Center line: median; box limits: upper and lower quartiles; whiskers: the most extreme data point which is not more than 1.5 times the interquartile range from the box. The *p*-values were computed using *t*-tests within linear models, adjusting for age, sex, dichotomized background staining level, and, for ROSMAP samples, post-mortem interval. **(E)** Correlation between the proportion of microglial CD74^high^ and the annual rates of cognitive decline in 59 ROSMAP individuals. The linear regression line is shown, with the shaded area representing the 95% confidence interval. The *p*-values were computed using *t*-tests within linear models, adjusting for age, sex, dichotomized background staining level, and post-mortem interval. Each dot represents an individual sample. **(F)** Estimate of cognitive slops are shown for two groups of individuals: the top 10 (red) and bottom 10 (blue) based on the proportion of CD74^high^ microglia. The slope was estimated by fitting annual cognitive scores of patients in each group with a linear mixed effect model with a random slope.

ROSMAP participants also undergo annual neuropsychologic evaluations while they are alive, allowing us to calculate person-specific slopes of cognitive decline^23^, and clinicians evaluated each participant for a diagnosis of AD dementia. Interestingly, the proportion of CD74^high^ microglia cells is significantly increased in individuals with a diagnosis of AD dementia (p=0.038) (**Fig. 4D**). Further, we find that individuals with a greater proportion of CD74^high^ cells have a significantly faster rate of cognitive decline (**Fig. 4E**), consistent with the greater prevalence of these cells in demented individuals. An alternative visualization of this trend presents the trajectories of cognitive performance of individuals who had the highest or lowest proportions of CD74^high^ microglia. We observed that individulals who have the highest proportion of CD74^high^ cells (red) have a faster cognitive decline than those have the lowest proportion of CD74^high^ cells (blue) (**Fig. 4F**). These clinical and cognitive measures were unfortunately not available in the NYBB samples.

The richness of these longitudinal data can be further modeled as rates of cognitive decline are not linear. Prior modeling of ROSMAP data has shown that there is an inflection point around the time that an individual begins to display mild cognitive impairment; after the inflection point (identified at the participant level), the rate of cognitive decline accelerates, and the participant enters the phase of terminal decline, which typically occurs in the last 3 years of life^24^. However, not all participants have entered this phase of decline at the time of death. We therefore leveraged this prior modeling of longitudinal cognitive data^23^, and we partitioned ROSMAP participants into those who died while in the terminal decline phase and those that had not entered this phase of the disease. When we compared the two groups, we found that these CD74^high^ cells are increased in frequency among individuals who are in the phase of terminal decline (p=0.015) compared to individuals who are in the earlier phase of cognitive decline (**Fig. 4F**). Overall, it seems that CD74^high^ microglia may play a role in the final stage of AD: in contributing to the loss of cognitive performance after the accumulation of the amyloid and tau proteinopathies that define AD.

### Morphological characterization of CD74^high^ microglia

We assessed the morphological characteristics of the CD74^high^ cells that we segmented in the ROSMAP and NYBB samples focusing on their roundedness. We evaluated the frequency of microglia classified as round among CD74^high^ microglia and among CD74^low^ microglia (**Fig. 5A**). In ROSMAP, the proportion of CD74^high^ microglia that are also rounded (median, 8%; Q1 – Q3, 4% – 10%) is significantly higher than in CD74^low^ microglia (median, 5%, Q1 – Q3, 2% – 7%) (p = 1.4 × 10^-6^), and we report the same association in the NYBB with median 5% (Q1 – Q3, 3% – 7%) of CD74^high^ microglia being rounded vs. median 2% (Q1 – Q3, 1% – 4%) of CD74^low^ microglia (p = 2 × 10^-13^). Thus, while CD74^high^ microglia are more likely to have the rounded morphology traditionally attributed to activated microglia, not all morphologically activated cells appear to be CD74^high^; hence, it is possible that there are molecularly distinct subtypes of microglia displaying an activated morphology. At this time, we therefore don’t have a molecular proxy for the classic stage III rounded microglia; rather, high CD74 expression and a rounded morphology may be markers of distinct microglial molecular programs that frequently co-occur in individual microglia.

**Fig. 5:**
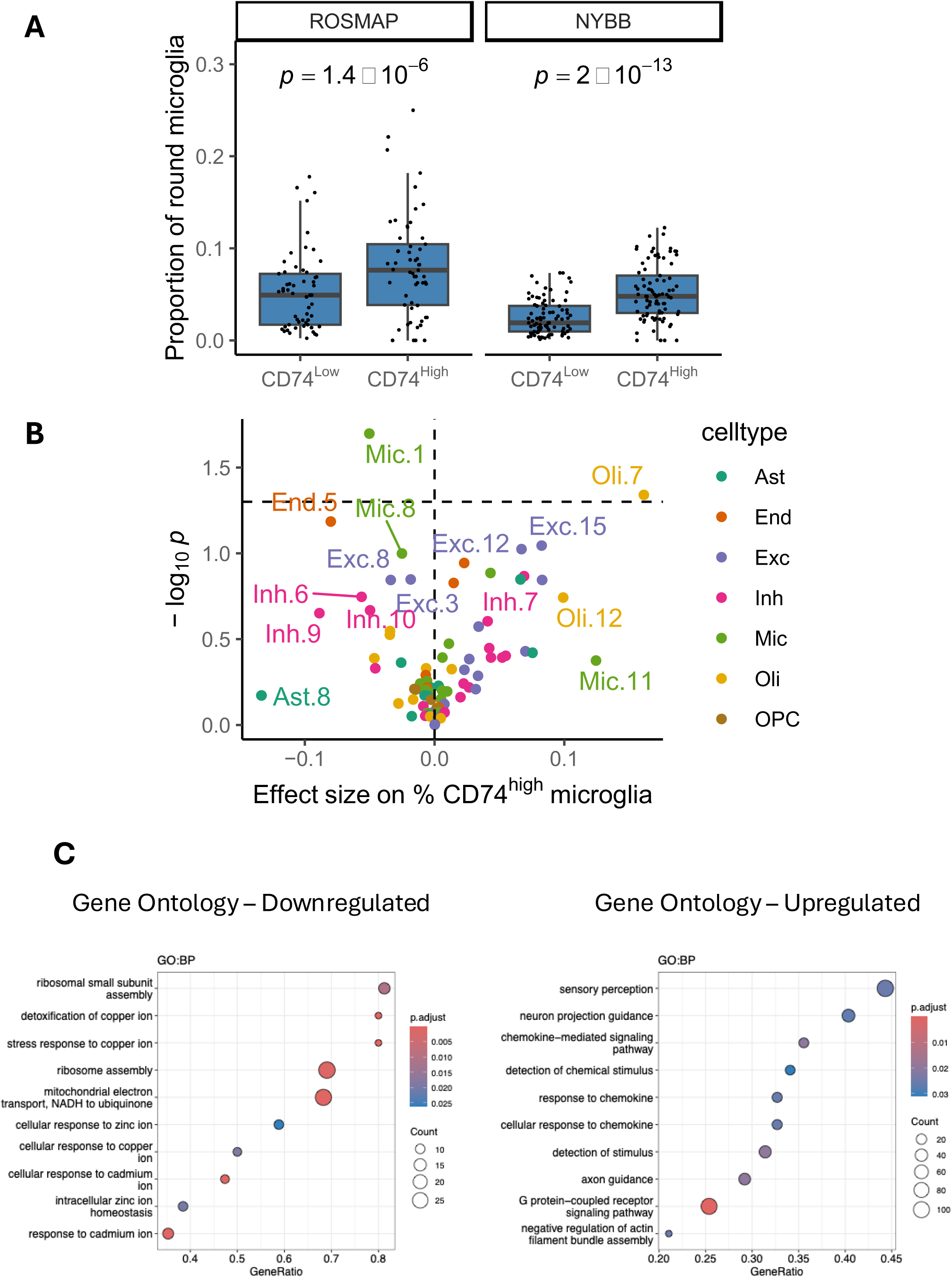
Analysis of CD74^high^ microglial morphology, associations with cell subtype distributions, and implicated pathways. **(A)** Box plots illustrating the association between the proportion of rounded microglia and CD74^high^ microglia in both cohorts. As each individual contributed one value to the CD74^high^ group and another to the CD74^low^ group, the *p*-values were computed with Wilcoxon signed rank tests for paired samples. Center line: median; box limits: upper and lower quartiles; whiskers: the most extreme data point which is not more than 1.5 times the interquartile range from the box. **(B)** Volcano plots showing the association between CD74^high^ microglia proportion and cell subtype propotions in our previous snRNA-seq study. Each point represents a specific cell subtype, colored by cell type: microglia (Mic), astrocytes (Ast), endothelial cells (End), excitatory neurons (Exc), inhibitory neurons (Inh), oligodendrocytes (Oli), and oligodendrocyte precursor cells (OPC). The effect sizes and *p*-values were computed using linear models adjusted for dichotomized background staining level. Dashed horizontal and vertical lines indicate *p* = 0.05 and effect size = 0, respectively. **(C)** Functional enrichment of differentially expressed genes in CD74^high^ microglia. Microglia from a glioblastoma sample was profiled using CITE-seq, which simultaneously measures transcriptome and surface protein levels at single-cell resolution. The CD74^high^ subset was defined based on surface CD74 expression levels. Differentially expressed genes between the CD74^high^ and CD74^low^ subsets were identified, and their Gene Ontology enrichment are displayed. (Left) Pathways enriched among genes positively associated with CD74^high^ cells, and (right) pathways enriched among genes negatively associated with CD74^high^ cells. Dot size represents the number of genes in each pathway, and color indicates adjusted *p*-value (*p*. adjust).

### Examination of the association between CD74^high^ microglia and specific microglial states

Over the past decade, brains derived from the ROSMAP cohort have been characterized using various molecular approaches. Our recent single-nucleus RNA-sequencing studies revealed complex glial cell substates in the human cortex, with some of these subtypes being associated with specific AD traits among 1,638,882 high-quality nuclei profiles from 437 participants^3^. Microglia were partitioned into 16 subpopulations while astrocytes were partitioned into 10 subpopulations^3^. In order to investigate whether CD74^high^ microglia are associated with a specific microglial or astrocyte subpopulation, a linear regression analysis of cell proportions was conducted (**Fig. 5B**).

The correlation between the proportion of CD74^high^ microglia and the proportion of different subtypes of microglia from the single-nucleus RNA sequencing data (n=38 ROSMAP participants) revealed a negative correlation between the proportion of Mic.1, which is a proliferative subpopulation, and CD74^high^ microglia calculated from our IF images (p=0.02). No significant correlation was observed with astrocyte subpopulation frequencies (**Fig. 5B**). Finally, since genotypes are available for the ROSMAP samples, we also assessed whether AD susceptibility variants influence the proportion of CD74^high^ cells. We assessed 83 SNPs tagging the susceptibility variants from a recent AD GWAS study^25^ and found that none of them is associated with the quantity of CD74^high^ cells at the level of FDR < 0.05.

### Transcriptomic and proteomic characteristics of CD74^high^ cell

To understand the transcriptional profile of CD74^high^ cells, we accessed recently produced CITE-seq data in which human microglia were isolated from fresh autopsy and surgical tissue by mechanical dissociation, followed by anti-CD11b magnetic bead purification, staining with a custom microglial antibody panel (**Supplementary Table. 1**) and single-cell RNA sequencing using the chromium platform (Methods)^26^. One library of magnetic-bead purified microglia from a glioblastoma sample had high cell viability and the best quality characteristics of all 9 generated libraries. Using this library, we leveraged the CD74 protein expression level from the CITEseq panel to classify the CD74^high^ microglia using the same measure of >2 standard deviations from the mean CD74 expression levels; as seen in **Supplementary** Fig. 4B, the CD74^high^ microglia cluster to the left of the graph, suggesting that they are quite different transcriptomically, along the UMAP 1 dimension, which captures the greatest amount of variance in these microglial transcriptomes. Using Gene Ontology Biological Process (GO:BP) enrichment analysis, we identified specific pathways that are upregulated in CD74^high^ cells included the G protein-coupled receptor signaling pathway (p=2.98×10^-5^) and chemokine-related pathways such as response to chemokine (p=0.026) and cellular response to chemokine (p=0.026) (**Fig. 5C, Supplementary Materials**). In contrast, pathways that were significantly downregulated in CD74^high^ cells included particularly those involved in the cellular response to zinc and copper ions (FDR<0.05) (**Fig. 5C, Supplementary Materials**). Specifically, the pathways response to copper ion (p = 0.037), cellular response to copper ion (p = 0.019), cellular response to zinc ion (p = 0.026), response to metal ion (p = 1.2 × 10^-5^), and stress response to copper ion (p = 2.44 × 10^-6^) were significantly downregulated in CD74^high^ cells. The main genes involved in these pathways included several metallothionein family members, such as *MT1M*, *MT2A*, *MT1X*, *MT3*, *MT1E*, and *MT1G*. Notably, the individual genes contributing to these pathways were not significantly differentially expressed at the RNA or protein level in CD74^high^ cells (**see Supplementary Materials**). The data suggest that CD74^high^ cells express lower levels of metallothionein genes, which play a key role in regulating intracellular zinc and copper levels by binding and sequestering free zinc and copper.

### Brain MIF protein expression, a ligand of CD74 increases with cognitive decline

Delving into the possible role of CD74^high^ microglia, one of the high-affinity ligands of CD74 is macrophage migration inhibitory factor (MIF), a pro-inflammatory cytokine that has been reported to contribute to cognitive impairment in individuals with AD dementia in several studies^27–29^. Using proteomics data from the DLPFC of the ROSMAP cohort, we evaluated the association between MIF expression and clinical AD diagnosis (n=864). MIF expression was significantly increased in individuals with a clinical diagnosis of AD dementia (p=0.02) compared to non-cognitively impaired individuals (**Fig. 6**), matching the increased proportion of CD74^high^ microglia observed in individuals with dementia. (**Fig. 4D**). MIF protein expression was also associated with the slope of cognitive decline (p= 0.0015). However, unlike the proportion of CD74^high^ microglia (**Fig. 4D**), MIF protein levels in our bulk cortical profile were also significantly increased in individuals with AD pathology (n=875)(p=0.011) (**Fig. 6**). These findings suggest that MIF upregulation may occur first, in relation to proteinopathy, contributing to an increase in CD74^high^ microglia which appear to primarily influence cognitive decline.

**Fig. 6:**
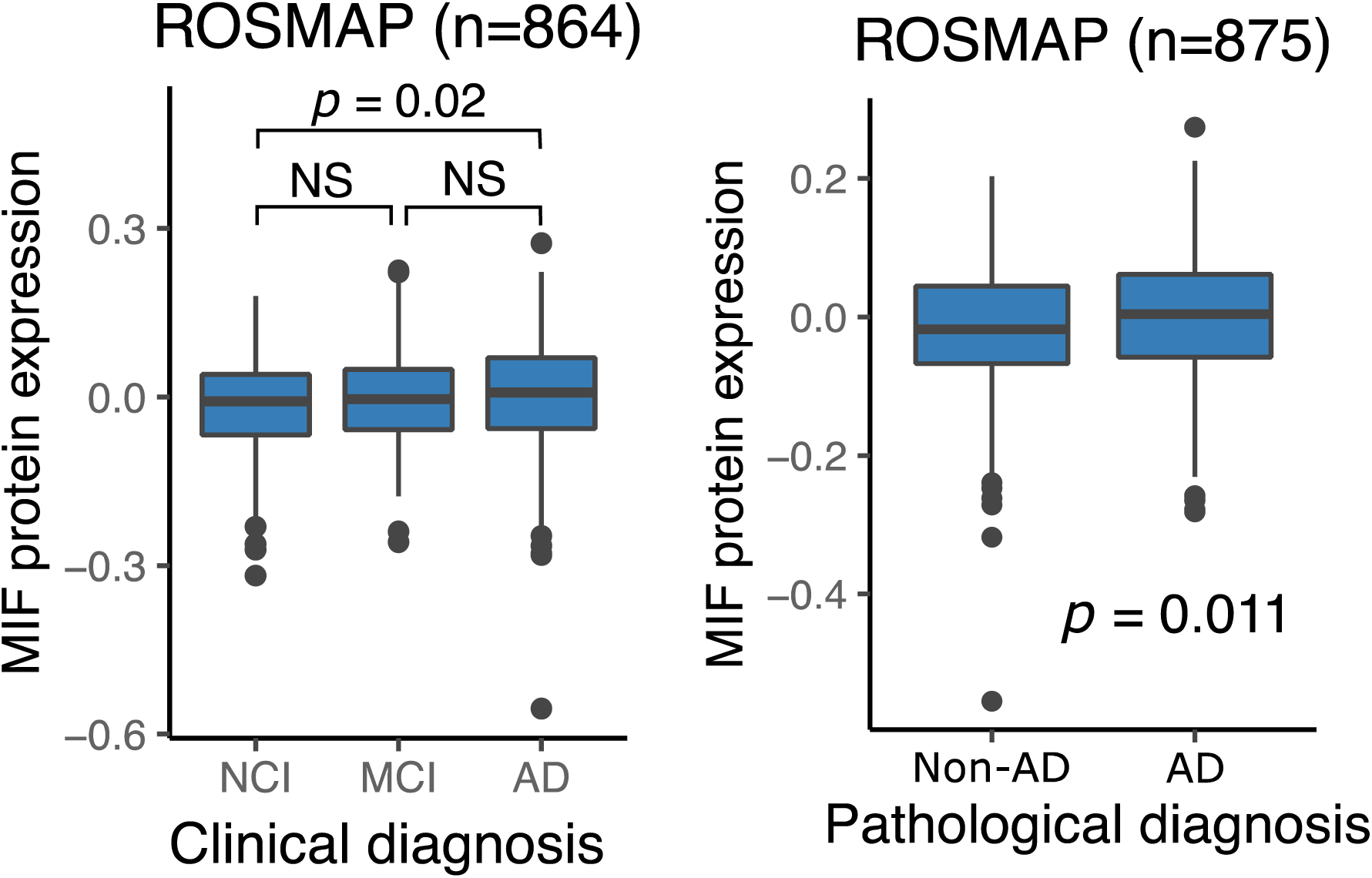
Association of MIF protein expression with AD clinical and pathological diagnoses. (Left) Boxplots show MIF protein expression levels in individuals with no cognitive impairment (NCI), mild cognitive impairment (MCI) and clinical AD. (Right) Boxplots show MIF protein expression levels in individuals with or without AD. The *p*- values were computed with Wilcoxon rank sum tests. Center line: median; box limits: upper and lower quartiles; whiskers: the most extreme data point which is not more than 1.5 times the inerquartile range from the box. The dots represent outliers.

## Discussion

In this study, we conducted an in-depth evaluation of a single marker protein, CD74, to illustrate the utility of our moderate-throughput image capture, segmentation, and analysis pipeline that is optimized for human microglia. Since most microglia express CD74, we used a rigorous mathematical approach to define the subset of microglia characterized by high expression of CD74 (**Fig. 4A and 4B**), facilitating reproducible and cross-study data collection. We test the generalization of our approach by characterizing brain sections from two very different brain collections (**Table. 1**): ROSMAP, prospectively collected brains from two cohort studies managed by a single set of investigators, and NYBB, a more typical brain bank accepting brain donations from individuals with different antemortem diagnoses. Both collections return the same results (**Fig. 4C**).

The subset of microglia with high levels of CD74 expression was initially studied in a smaller number of samples and demonstrated an association with AD^2^. This initial finding served as the motivation to conduct a more extensive evaluation of the strategy for defining CD74^high^ cells; we now have obtained robust results suggesting that the frequency of the CD74^high^ subset of microglia is not associated with the accumulation of AD-related amyloid and tau proteinopathies (**Fig. 4C**). However, it is associated with cognitive impairment and AD dementia (**Fig. 4D**), particularly in the phase of terminal decline (**Fig. 4F**). These observations suggest that this particular microglial subtype might be involved in a later stage of the cascade of events leading to cognitive decline.

To explore a possible molecular mechanism by which CD74^high^ microglia may contribute to cognitive decline, we investigated the association between MIF protein expression and both clinical and pathological AD diagnoses. Interestingly, MIF, a high-affinity ligand of CD74, has been reported to contribute to cognitive impairment in individuals with AD dementia in several studies^27–29^. Specifically, higher MIF expression has been detected in the CSF of AD patients^28,29^ compared to control patients, and higher MIF levels were associated with accelerated cognitive decline in MCI and mild dementia^28^. Our findings are consistent with previous studies, showing that increased MIF protein expression in the DLPFC of individuals with dementia (**Fig. 6**). This provides additional support for the potential role of the CD74^high^ microglial subset in cognitive impairment via a higher level of MIF-CD74 signaling. The examination of the transcriptomic and proteomic characteristics of purified CD74^high^ microglia reveals upregulation of genes involved in chemokine response and downregulation of metallothionein genes involved in pathways related to cellular responses to metal, copper, and zinc ions, as well as the stress response to copper ions (**Fig. 5C**). Metallothioneins genes are metal-binding proteins involved in metal ion regulation ^30^ and play a central role in sequestering zinc and copper to prevent toxicity. An enrichment of copper and zinc within amyloid plaques has been reported ^31,32^. Therefore, the downregulation of metallothionein genes may have significant consequences on amyloid burden and clearance, as studies have shown that metallothioneins can prevent copper-induced aggregation of amyloid-β peptides^33^. Therefore, the downregulation of pathways related to cellular responses to metal, copper, and zinc ions in CD74^high^ cells may contribute to the accumulation of amyloid plaques in individuals with AD, potentially leading to the cognitive decline observed (**Fig. 4C**). However, this hypothesis requires further validation.

Our automated image segmentation approach also enabled us to assess the function of these cells through an analysis of morphological features. High levels of CD74 identifies a microglial subtype that is significantly more rounded (fewer processes, lower compactness value) than other microglia in both sets of brain samples that we characterized (**Fig. 3C**). Compactness is a key morphological feature for classifying microglia in neuropathological studies; stage III microglia are considered to be activated and are characterized by thick, retracted processes and a rounded shape^34^. Previous studies reported that, in the neocortex, the proportion of stage III microglia are associated with AD traits in ROSMAP participants^20,35,36^. However, those studies rely on manual segmentation, a method known to have significant limitations with inter-rater variability when counting and classifying microglia into three distinct morphological stages. The challenges associated with the reliability of manual segmentation served as a key motivation for developing an automated segmentation approach on an entire tissue section in this study. This approach facilitated the morphological evaluation of microglia and enabled the profiling of a large number of microglia in each tissue section, enhancing our statistical power. We also developed an operational definition for microglia with a low compactness score independent of CD74 staining, which are associated with both proteinopathies and cognitive decline, similar to what has been reported for stage III microglia^20^. Thus, our automatically segmented small, compact microglia appear to be consistent with manually segmented stage III microglia: both cell subtypes have the same properties in relation to AD-related traits. CD74^high^ microglia, while more compact than other microglia, are only associated with cognition-related outcomes. While a modest effect on the pathologic outcomes cannot be excluded at this time given our moderate sample size, the magnitude of such an effect would be much weaker than that of stage III/small rounded microglia. Thus, CD74 staining helps us to resolve a distinct subset of microglia which is involved at the late, symptomatic phase of aging-related cognitive decline. These results are consistent with single cell studies of human microglia which implicate a microglial subtype enriched for the DAM2 RNA signature in amyloid and tau proteinopathy^3,4^. These DAM2^high^ cells are also small and rounded^4^, and they are enriched in the vicinity of neuritic amyloid plaques^37^. In these single cell/nucleus data, CD74^high^ microglia are a distinct subset that have some enrichment for DAM1 genes and are not associated with AD proteinopathies. Therefore, thanks to CD74, we begin to resolve subtypes of microglia that have an activated morphology (classical stage III microglia) into at least two subtypes which have distinct associations with AD-related outcomes and likely different functions. This is consistent with their partially overlapping but distinct transcriptomes in sc/snuc studies^4^.

Larger, better powered studies and investigation with additional markers associated with the CD74^high^ microglial subset are needed to better understand the mechanisms by which this particular subset contributes to cognitive impairment in AD patients. CD74^high^ microglia could provide a key target for therapeutic development as they appear to be closely connected with the most important outcome in AD: cognition.

## Acknowledgements

We thank the participants in the ROS and MAP studies as well as donors in the NYBB for their generous contribution to the study of cognitive aging and AD. The ROSMAP studies are supported by several grants from the National Institutes of Health: P30AG10161, P30AG72975, R01AG15819, R01AG17917. U01AG46152, U01AG61356. The ROSMAP resources can be requiested at www.radc.rush.edu and www.synapse.org. The NYBB is supported by the Columbia University Irving Medical Center AD Research Center grant, P30AGO66462. Data presented in this manuscript were generated and analyzed through support by the Chan-Zuckerberg Initiative’s Neurodegeneration Challenge Network grant CS-02018-191971 as well as grants R01AG070438, U01AG061356, RF1AG057473, and R01AG048015.

---

**Supplementary Fig.1:**
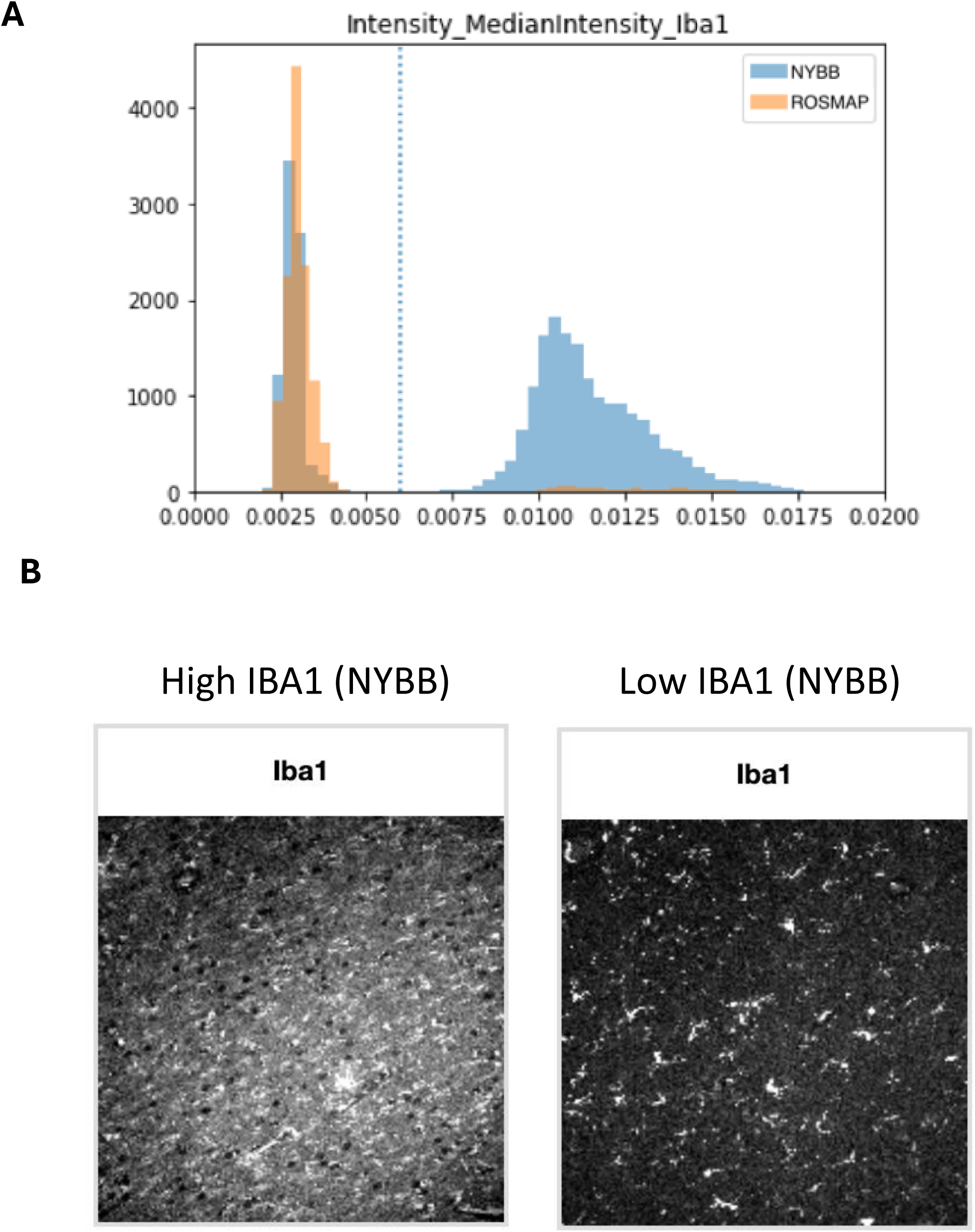
Dichotomy in the background staining level between cohorts and across samples. **(A)** Histogram illustrating variation in IBA1 channel background intensi’es across image ’les in two cohorts (ROSMAP: orange, NYBB: blue). The median pixel intensity of each image tile was used as a proxy for background intensity, as microglial cells occupy only a small fraction of the total pixels. **(B)** Representative IBA1- stained images (white) from samples with high (left) and low (right) background staining levels within the same cohort.

**Supplementary Fig.2:**
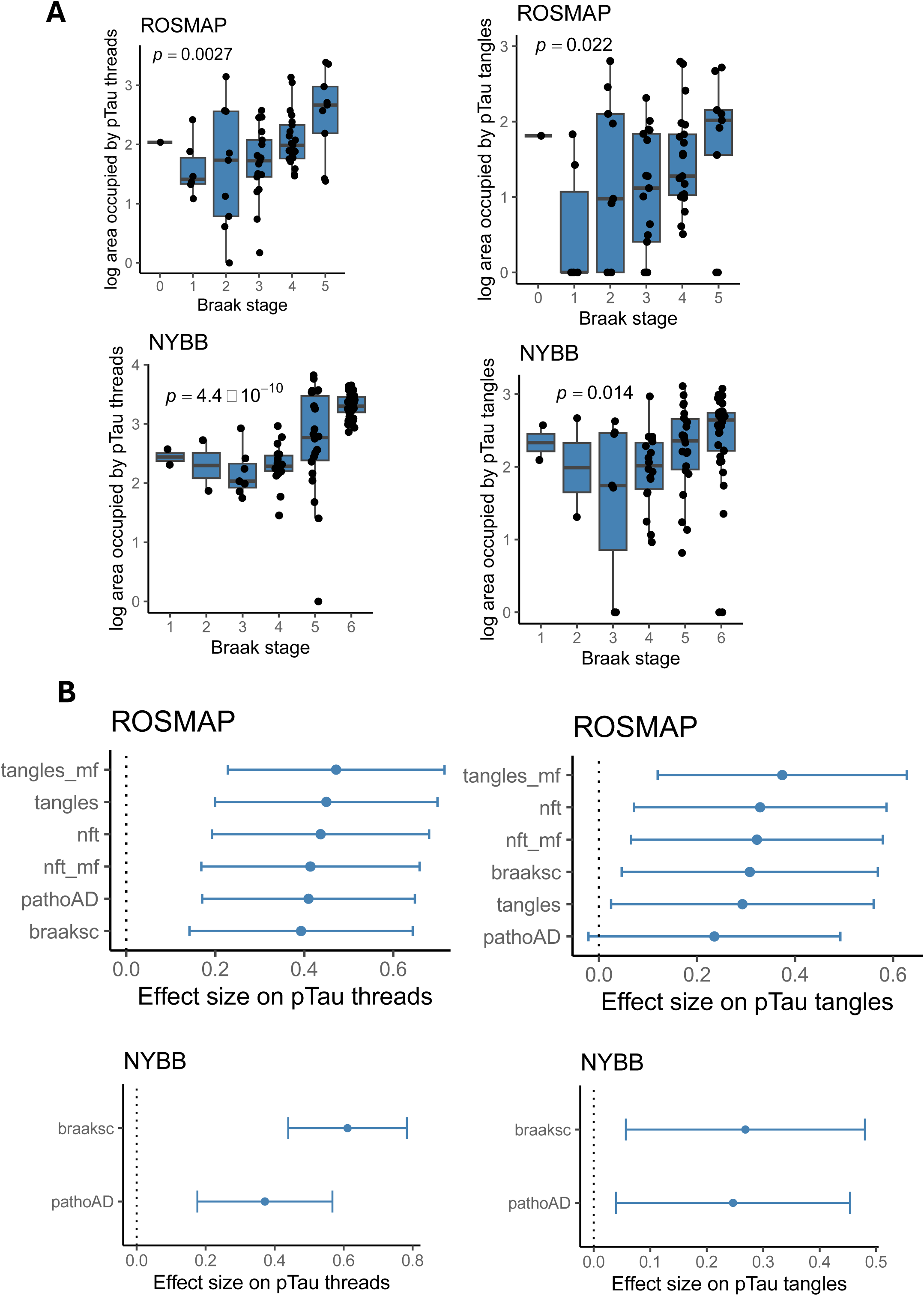
Association between Tau pathology staining, Braak stage, and AD pathological traits. **(A)** Box plots showing the association between the area occupied by pTau threads (left) and tangles (right) and Braak stage in the ROSMAP and NYBB cohorts. Each dot represents an individual sample. Center line: median; box limits: upper and lower quartiles; whiskers: the most extreme data point which is not more than 1.5 times the interquartile range from the box. **(B)** Forest plots show the effect size of each pathological trait (Braak stage and AD pathologies) on IF-based pTau object area (pTau threads and tangles) in a linear model adjusted for age, sex, and dichotomized background staining level. Post-mortem interval was also added as a covariate for ROSMAP samples. The error bars represent 95% confidence intervals.

**Supplementary Fig.3:**
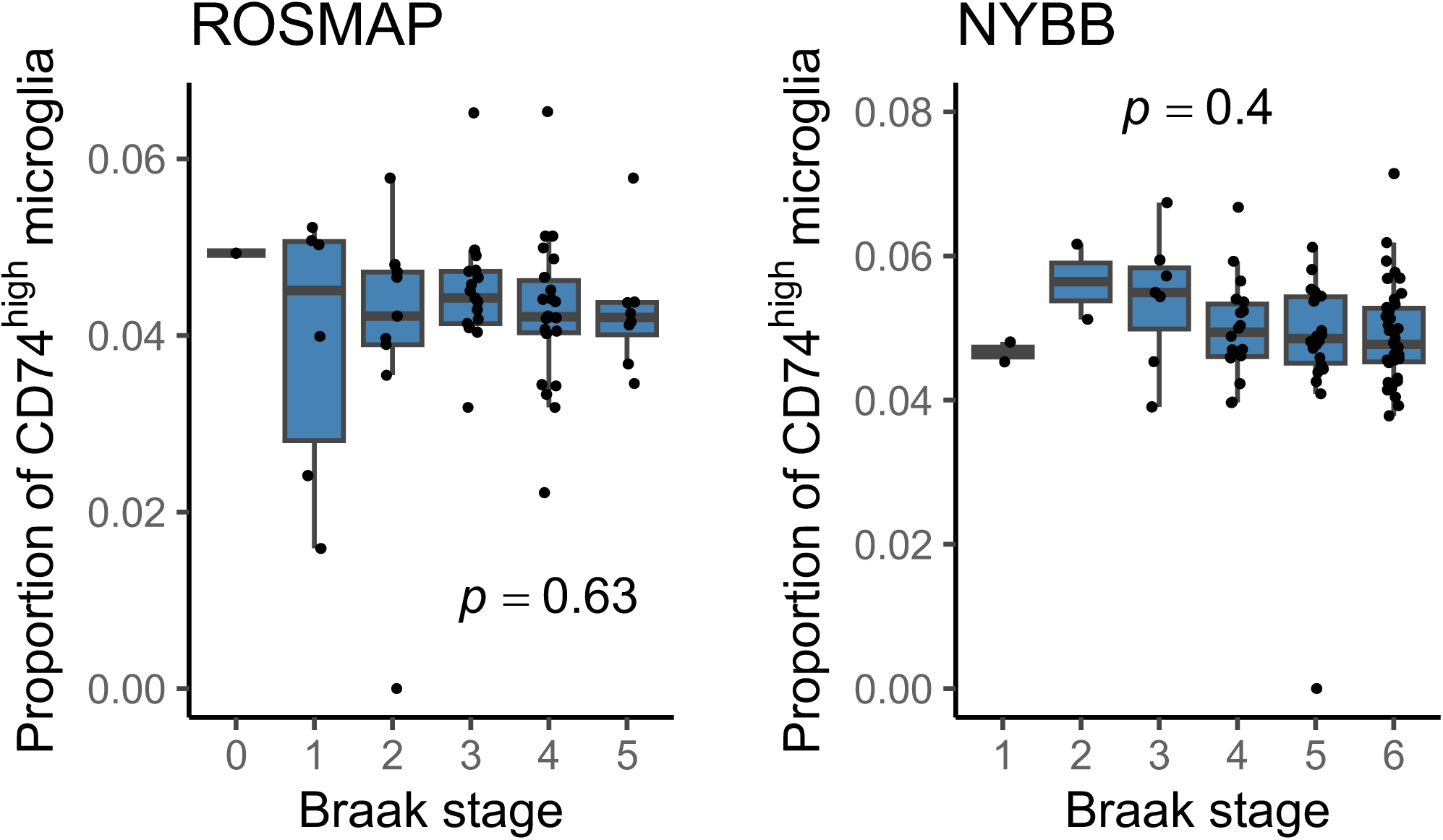
Association between CD74^high^ microglia and Braak stages. The boxplots show the associations between the proportions of CD74^high^ microglia and Braak stages in ROSMAP (left) and NYBB (right) cohorts. Center line: median; box limits: upper and lower quartiles; whiskers: the most extreme data point which is not more than 1.5 times the interquartile range from the box. Each dot represents an individual sample.

**Supplementary Fig. 4:**
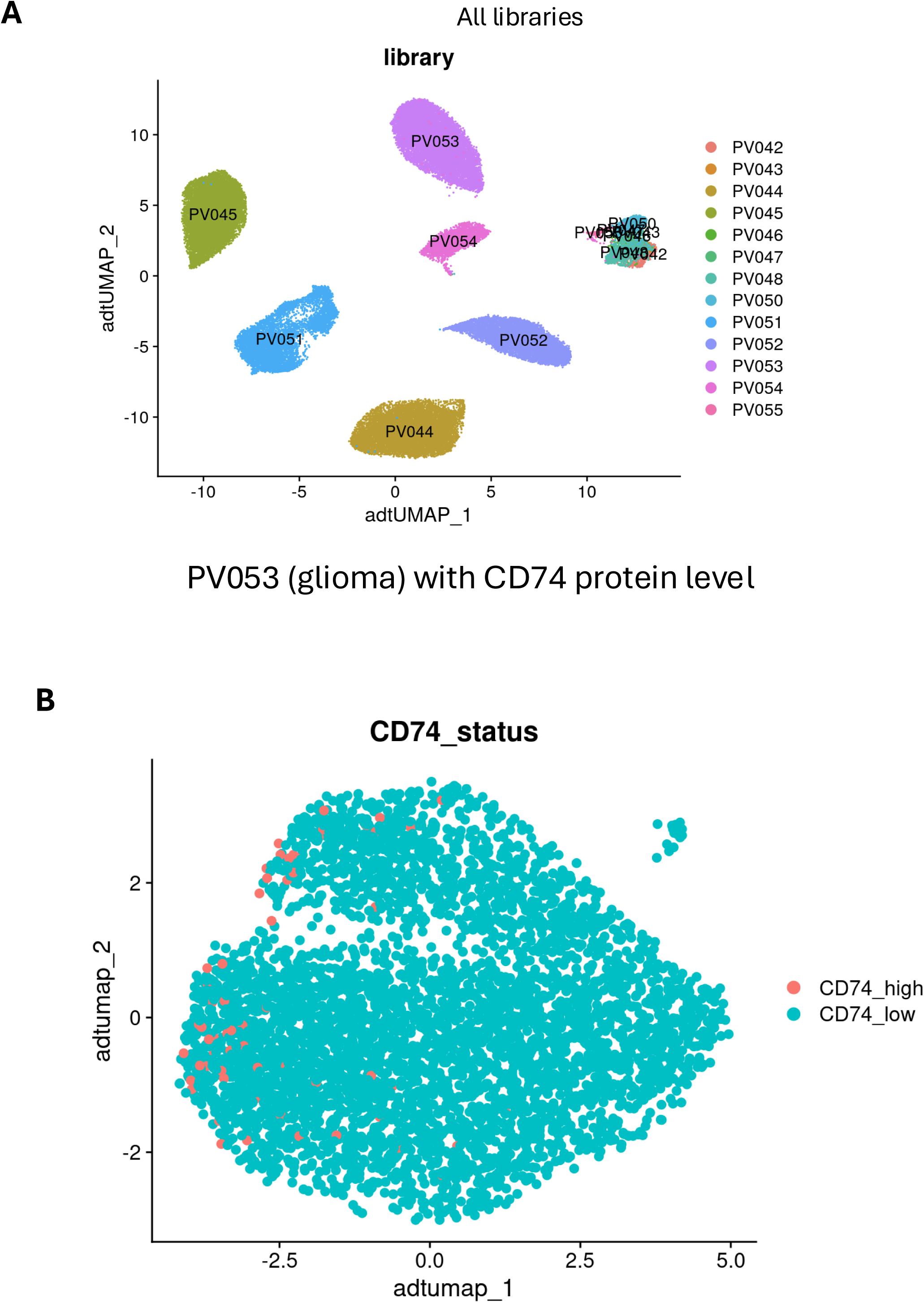
CD74^high^ cells in microglia CITE-seq dataset. (**A**) UMAP dimentionality reduction of protein abundance in 13 CITE-seq libraries of microglia (PV042-PV055). (**B**) Protein-based UMAP of microglia from a glioma sample (PV053). In red, microglia with high CD74 abundance; in blue, microglia with low CD74 abundance.

**Supplementary Table. 1:**
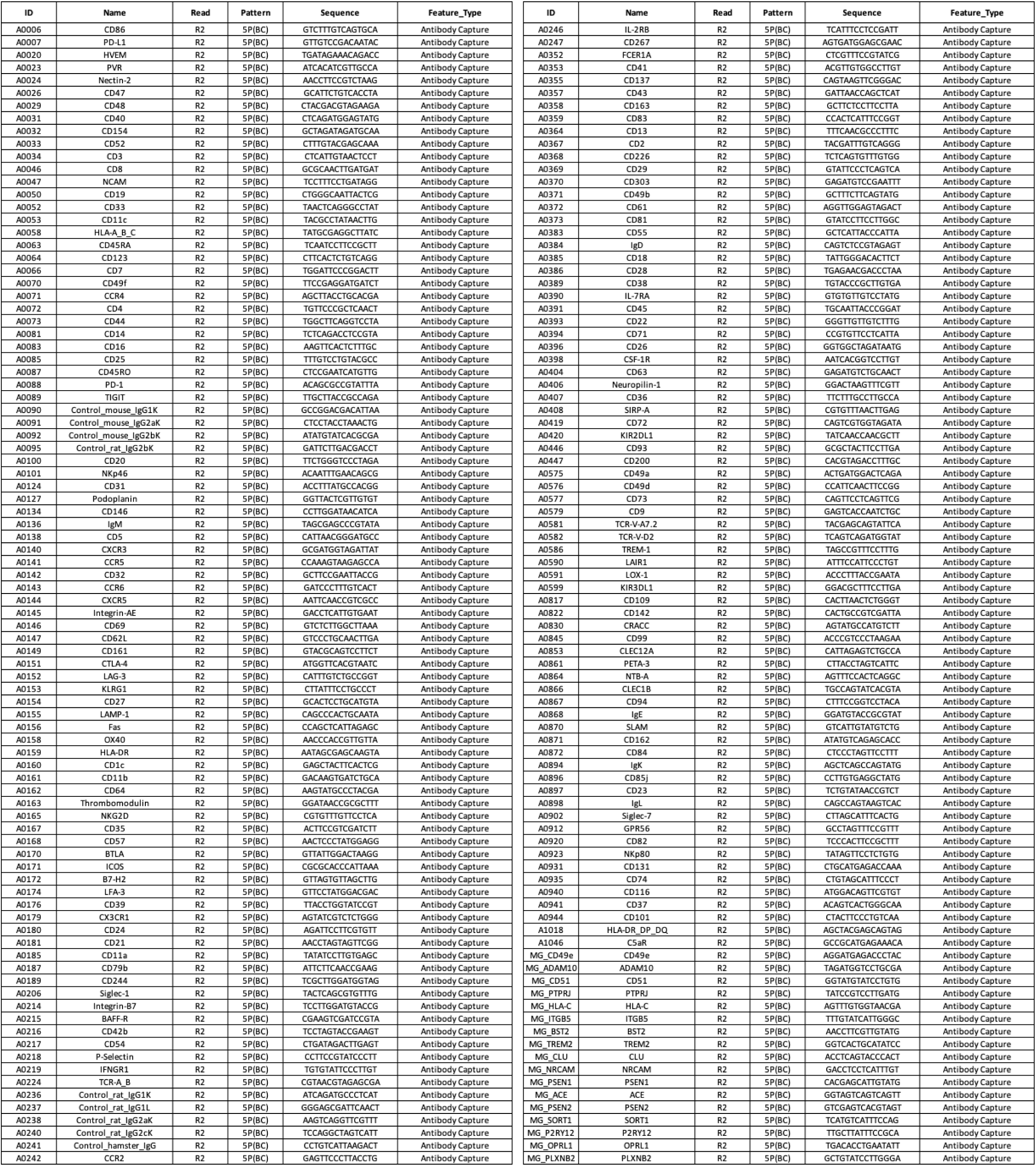
List of custom microglial antibody panel used for CITE-seq profiling of microglia from glioblastoma samples.

